# Mechanics of wood boring by beetle mandibles

**DOI:** 10.1101/568089

**Authors:** Lakshminath Kundanati, Nimesh R. Chahare, Siddhartha Jaddivada, Abhijith G. Karkisaval, Nicola M. Pugno, Namrata Gundiah

## Abstract

Wood boring is a feature of several insect species and is a major cause of severe and irreparable damage to trees. Adult females typically deposit their eggs on the stem surface under bark scales. The emerging hatchlings live within hard wood during their larval phase, avoiding possible predation, whilst continually boring and tunneling through wood until pupation. A study of wood boring by insects offers unique insights into the bioengineering principles that drive evolutionary adaptations. We show that larval mandibles of the coffee wood stem borer beetle (*Xylotrechus quadripes*: Cerambycidae) have a highly sharp cusp edge to initiate fractures in *Arabica* wood and a suitable shape to generate small wood chips that are suitable for digestion. Cuticle hardness at the tip is significantly enhanced through zinc-enrichment. Finite element model of the mandible, based on the differential material properties at the tip, intermediate and base regions of the mandible, showed highest stresses in the tip region; these decreased to significantly lower values at the start of the hollow section. A hollow architecture significantly reduces bending stresses at mandibular base without compromising the structural integrity. A scaling model based on a fracture mechanics framework shows the importance of the mandible shape in generating optimal chip sizes. These findings contain general principles in tool design and put in focus interactions of insects and their woody hosts.

## 1. Introduction

Wood-boring (xylophagous) beetles cause irreversible damage to forests, crops and timber due to their impact on transport of sap and nutrients in stems [1, 2]. The larval ability to tunnel inside wood allows them to successfully evade their natural enemies when accessing a nutritional source for which there is reduced competition. The remarkable success of xylophagous insects mainly lies in the larval ability to repeatedly cut through hard substrates over long duration. Although these insects are among the leading threats to forests and tree plantations across the world, little is known about the mechanics of their mandibles in wood boring. Mandibular design requires a structure which can be repeatedly used to easily cut through wood fibrils without itself undergoing fracture, wear or failure. Another essential requirement is the generation of small wood chip which may easily be ingested by the insect. Of equal importance is the toughness of the mandibular cuticle and its ability to withstand the structural stresses during boring. A combination of toughness and hardness facilitates diverse functions in beetle mandibles such as cutting, digging, combat etc [3, 4].

The cuticle of insect mandibles is mainly composed of proteins, lipids, and carbohydrates which are hierarchically organized into stratified layers to produce one of the toughest known natural composites [4, 5]. Mechanical properties of the cuticle can range from being hard in many larval or adult mouthparts to highly flexible in the intersegmental membranes; such adaptations permit a diverse functional repertoire of the cuticle in insects [6, 7]. The vast range of mechanical properties of the cuticle is dictated by sharp variations in its material composition through incorporation of transition metals and halogens [4, 8, 9], or through *sclerotization* in which the cuticular proteins are stabilized through a variable degree of quinone reactions [10, 11]. Unlike the hard enamel coatings of mammalian teeth, hardness in insect mandibles is enhanced through metal-ion mediated crosslinking of chitin-protein complex molecules [12]. Such hardening mechanisms have been reported in diverse insect mandibles of termites, leaf cutter ants, and jaws of a nereid worm, *Nereis virens*, etc. through incorporation of transition metals like zinc and manganese [13–17]. To investigate the mechanisms of wood boring in insects, we studied the coffee white stem borer (CWSB), *Xylotrechus quadripes*, which bores through coffee stems (*Coffea Arabica* L.) during an 8-10 month long developmental period [18]. The beetle larvae chew through the stem, thereby severely reducing the production of coffee beans, and ultimately cause the destruction of entire plantations [2].

The goals of our study are to characterize the structure-property relationships of the CWSB larval mandibles and analyze its response to the mechanical stresses encountered during wood chipping. First, we characterize the surface morphology and the compositional variation at different locations in the mandible using scanning electron microscopy with X-ray microanalysis. We also quantify the three-dimensional structure of the mandible using microtomography. Second, we estimate the mechanical properties of the mandibles at different locations using nanoindentation and correlate these with the compositional analysis. Third, we quantify the material properties of the wood substrate and compare these with that of the larval mandibles. Finally, we use a scaling model of two indenters cutting through a quasi-brittle wood material to characterize the relationship between the mandible shape and the chip sizes generated during cutting. The study explores biomechanical design principles and puts into focus the interactions of insects and their wood-hosts, and provides fascinating principles for engineers who design boring and cutting tools.

## 2. Materials and Methods

### 2.1. Insect model system

Fourth and fifth stage larvae of coffee white stem borer (*Xylotrechus quadripes* Chevrolat; Coleoptera, Cerambycidae; CWSB) were collected from uprooted infested coffee stems (*Coffea arabica* L.) from plantations in the Western Ghats in Karnataka, India, stored in 100% ethanol, and transported to the laboratory (Fig. 1A). Mandible specimens were dissected from the larvae, cleaned using ultrasonication, and prepared for the experiments (Fig. 1B). Once dissected, the mandibles of CWSB larvae were cleared of the muscles and other soft tissues, and imaged using bright-field microscopy. Specimens were embedded in acrylic resin and polished to achieve a flat surface for nanoindentation and compositional analysis.

**Figure 1.**
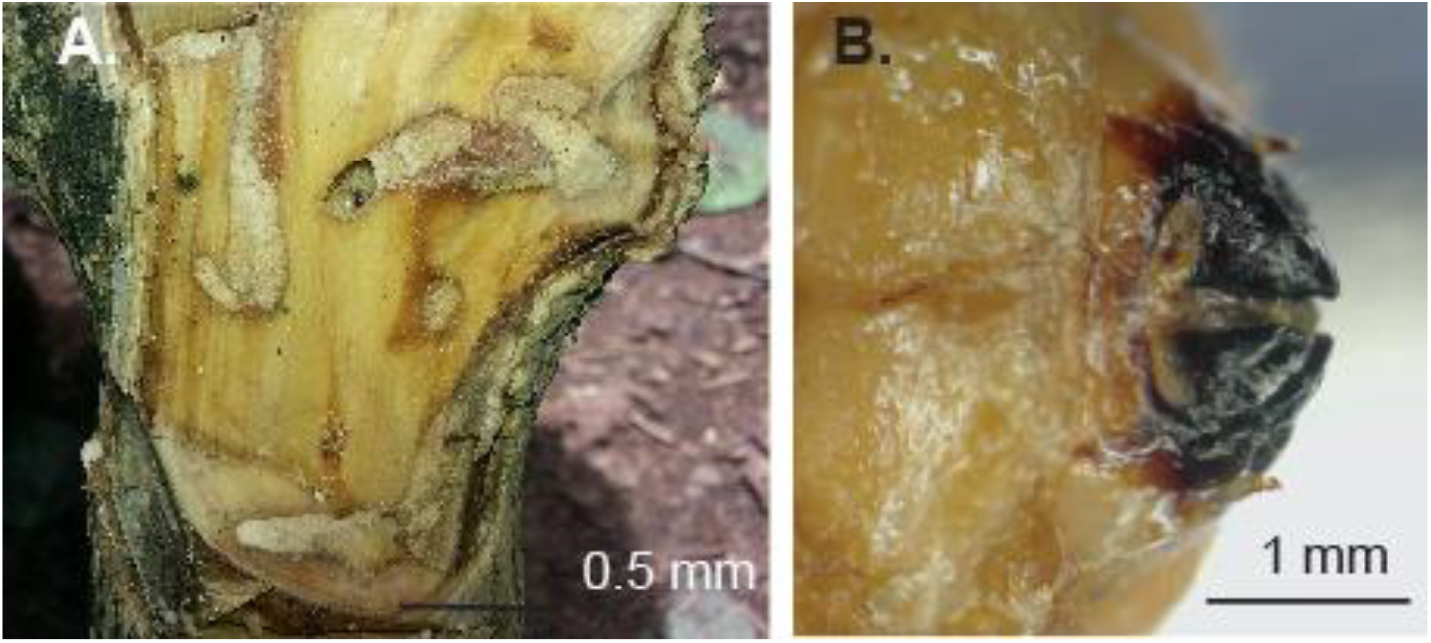
**A:** Transverse section of infested coffee wood shows damage to the wood by the Coffee Wood Stem Borer beetle (*Xylotrechus quadripes*: Cerambycidae; CWSB) larvae. **B:** Mandibles of the larvae sample highlights the sclerotized cutting regions of the larval body.

### 2.2. Scanning Electron Microscopy (SEM) and Energy Dispersive X-ray Spectroscopy (EDX)

The prepared mandibles were carefully mounted on double-sided carbon tape (Electron Microscopy Sciences, USA), attached to aluminum stub, and placed in a desiccator to avoid contamination. Specimens were transferred to a sputter coater (Bal-Tec SCD 500, Liechtenstein), the chamber purged thrice with argon, and the mandibles coated with a thin gold layer. A scanning electron microscope (Quanta 200 ESEM, FEI, The Netherlands) was used to image the samples with accelerating voltages between 5 and 20 kV. To quantify the elemental composition of mandibles, polished specimens were mounted on carbon tape and coated with 10 nm layer of gold. EDX detector in the SEM was used to scan the samples and the spectra collected using an accelerating voltage of 20 kV. A custom software (Genesis, FEI) was used to identify the individual peaks in a given spectrum. Variations in zinc along the length of the mandible were determined using a line scan. A pure zinc sample was used as reference for calibration of these spectra.

### 2.3. X-Ray microtomography

We used a non-destructive X-ray microtomography (microCT) technique to obtain three dimensional images of the mandible and its connecting musculature. To prepare the samples for microCT, the larval head containing the mandible was cleaned with ethanol to remove possible debris, and the labrum and clypeus were mechanically dissected from the body. The specimen was fixed in Bouin’s solution for 16 hours, dehydrated using graded solutions of ethanol (30% - 100%; 10 minutes each), and stained using iodine (1g in 10 ml of 100% ethanol) for 24 hours to improve the optical contrast between the cuticle and soft tissues [19]. A toothpick was used to mount the air-dried sample on the specimen holder in the microCT instrument (Xradia Versa XRM 500). A beam of X-rays of 60 kV accelerating voltage (6W) was incident on the sample and the projected plane was recorded to obtain a two-dimensional image. A set of 2500 projections was obtained by rotating the sample at small discrete angles about the vertical axis and a 3D structure was reconstructed using XRM software. A raw threshold was used to reduce the background noise for better visualization in the images. The reconstructed dataset was analyzed using AMIRA (v6.0.1, XEI Software, The Netherlands) to extract structural details including the connecting musculature and volume of the hollow regions within the mandible.

### 2.4. Uniaxial compression of wood samples

We used a uniaxial testing machine (Mecmesin Multitest 10-i, UK), equipped with a 10 kN load cell, to quantify the monotonic compression results from wood specimens (*Coffea arabica* L.). Specimens were prepared according to Indian Standard for wood testing (IS:1708) by machining sections in the parallel (2 cm × 2 cm × 8 cm; n=4) and in the perpendicular to the wood grain (2 cm × 2 cm × 10 cm; n=4). Fresh samples from recently harvested stems were machined in the specified dimensions, the specimen thickness measured using calipers, and the samples were mounted using compression platens in the testing frame. The sample was preloaded to 500 N and tested under compression (1 mm/sec) to characterize the material properties. Forces, recorded using a load transducer, were converted to engineering stresses using initial specimen dimensions, and the strains were calculated by measuring encoder displacements during specimen loading. Young’s modulus, calculated using a linear elasticity framework, was quantified using a secant modulus method using the initial linear region of the stress-strain curve that represents the elastic deformation regime.

### 2.5. Microindentation of wood samples

We also used microindentation experiments to quantify the hardness of the Arabica wood samples. Specimens were polished using a series of 400, 800, 1200, 2000 and 4000 grade sand paper, cleaned for a debris free surface, and indented in a CSM micro indenter using a standard Vickers tip with a maximum of 10 N load and at a loading rate of 5 N/min.

### 2.6. Nanoindentation

Ultrasonicated and cleaned CWSB mandibles, mounted in acrylic resin, were polished using sand paper with varying grit sizes (200, 400, 600, 800, 1000, 1200, 1500, 2000) and diamond paste to achieve flat surfaces for mechanical studies. We used nanoindentation experiments on the prepared specimens using a 3-sided pyramidal Berkovich indenter (tip radius = 150 nm) to characterize the local mechanical properties. The specimen was loaded at 50 μNs^-1^ up to a maximum load of 500 μN using a nanoindenter (TI 900-Tribo Indentor, Hysitron, Minneapolis, MN, USA). To reduce any possible effects of creep during unloading, we used a trapezoidal cycle consisting of 10 second loading, 5 second hold, followed by a 10 second unload period [20]. The resulting load-displacement data were analyzed using custom software to determine the contact stiffness (S), reduced modulus (E_R_), and hardness (H) based on the well-established Oliver and Pharr method [21]. The maximum displacement (h_max_) at peak load (P_max_) and S were determined using the experimental data corresponding to the unloading curve. Specifically, S was defined as slope of the upper portion of the unloading curve during initial stages of unloading. We used the standard relation from Oliver-Pharr method to relate the measured stiffness to the reduced modulus given by

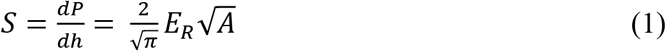

where A is the projected area of the contact between the indenter and the specimen. E_R_ is the reduced modulus defined using known parameters based on the modulus (E_i_) and Poisson’s ratio (ν_i_) of the indenter. Thus,

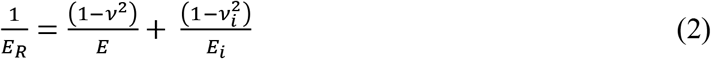

The material hardness, H, is defined as

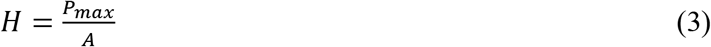

Where *P_max_* is the maximum load. A total of nine mandibles were tested with 8 indentations at each region separately. Statistical analyses to test for differences in the different regions were performed using ANOVA with Bonferroni comparison using *multcompare* function in MATLAB. A p value of <0.05 was considered statistically significant.

### 2.7. Finite Element simulations

Individual microCT images were used to obtain a three dimensional model of the mandible. A surface reconstruction of the mandible boundary from each transverse section was obtained by segmenting the image using MATLAB R2014b (Version 8.4) (MathWorks, Natick, MA, USA). Stacks of boundaries, corresponding to sections from the base to the mandible tip, were smoothed using B-splines, and the neighboring splines joined together using a loft feature (SolidWorks 2013, Concord, MA, USA) to obtain a three dimensional model (Fig. 2A-B). The model was meshed using10-node tetrahedral elements (Hypermesh 9, Altair Hyperworks, Troy, MI, USA). A total of 16328 elements were initially generated to create a coarse mesh (length 20 μm) of the structure. We used the numerical finite element method (FE) to simulate a quasi-static condition of the mandible tip in contact with the substrate and loaded to simulate wood cutting. The structure was divided into three regions (Fig. 2C) with Young’s moduli values varying from 6.6 GPa in the tip region located ~15 μm from the mandible apex, 6.2 GPa in the intermediate region, and 6.13 GPa in the base region of the mandible; these were based on results from nanoindentation studies. We assumed the value of ν as 0.3. Each of the two mandibles move due to the action of adductor and abductor muscles that are located at the mandible base (Movie S1; Movie S2). The pivot axis for mandible rotation was assumed to be along the axis M-M’ which was obtained from the videography of the borer (Fig. 2D; Movie S1). Specifically, the individual mandible edges during movement were tracked and the lever axis was determined to be located near the abductor muscles. The presence of a pivot axis, located towards the abductor muscle attachment region, has also been reported in other model systems such as the larvae of weevils, wood-boring ants, and dragonflies [22, 23]. Points of attachment of the abductor and adductor muscles to the mandibular base were determined using image segmentation from X-ray microtomography (Movie S2). Three different boundary conditions were used to assess stress distributions on the mandible; first, a fully pinned case where nodes corresponding to the mandible base had zero translational displacements. Second, a semi pinned case in which nodes were fixed only at adductor (S1 surface) and abductor muscle (S2 surface) attachments located at the base region of the mandible. Finally, a pivot case was simulated using M-M’ as the lever axis. In this method, the movement of nodes in the mandible base, located in proximity to M-M’ axis, were constrained using rigid links to achieve the desired pivot motion. Uniform tractions of 10 MPa were applied at surface S1 to simulate the action of adductor muscle. Surface tractions of 35 MPa, based on published reports of rupture stress of Balsam wood, were applied to elements located at the mandible tip (~15 μm) in a direction normal to the surface in contact with the substrate [22]. Mechanical equilibrium equations for the discretized structure were solved at each node using the prescribed boundary conditions using a commercial solver (Abaqus 6.11, Dassault Systèmes, Providence, RI, USA). Displacement and stress distributions were quantified in various regions of the mandible; solution convergence was assessed by mesh refinement to within 5% error between consecutive meshes. The final mesh used in reporting the solutions had 134707 elements which had varying lengths from the tip to the mandible base.

**Figure 2.**
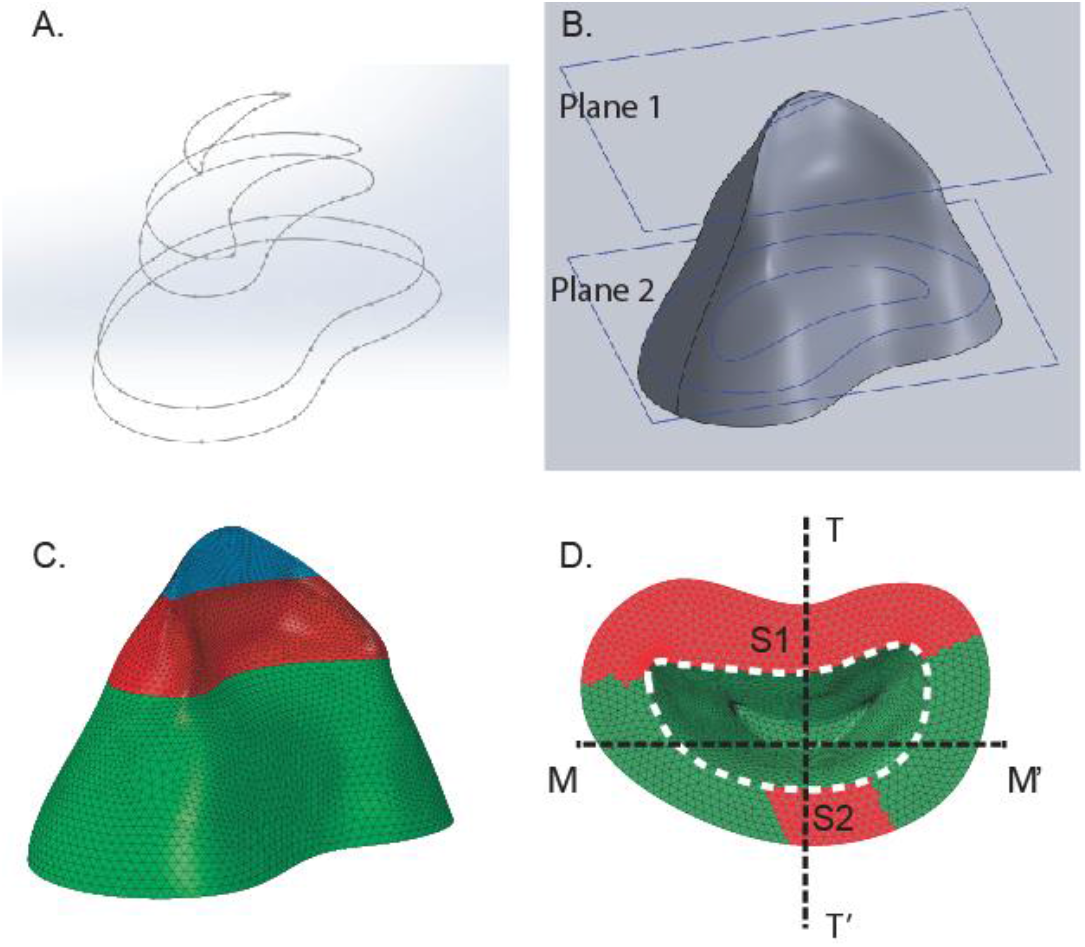
**A:** Splines were constructed using the microCT results at different heights from the base to the tip of the mandible that were used for constructing a 3D geometrical model **B:** The model after reconstruction is shown with cross sectional planes in the tip region corresponding to plane 1 (solid) and a hollow region (plane 2). **C:** The structure was meshed and divided into the tip (blue), intermediate (red), and base (green) regions of the mandible. **D:** Surfaces S1 and S2, located at the base of the mandible, corresponded to attachments of the adductor and abductor muscles. These, along with the axis of rotation (M-M’), were used to impose different boundary conditions to compare stress distributions on the mandible. T-T’ refers to the sagittal plane through the mandible.

### 2.8. Nanoindentation Scaling model of the mandibles in generating wood chips

Fracture toughness, *K_Ic_*, of the substrate during indentation is expressed in terms of the Young’s modulus, E, substrate hardness, H and the load for crack formation, P_max_, using the Lawn-Evans-Marshall model given by [24, 25]:

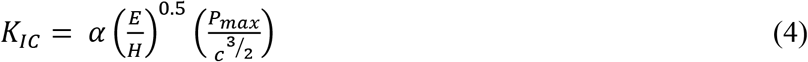

Where c is the length of the longest crack taken from the center of the indenter, and α is a dimensionless constant. For example, α=0.016 for Vickers indenter interacting with brittle materials. The maximum size chip formation occurs under an optimal condition given by:

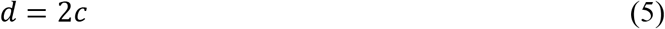

Where d is the separation distance between the indenters (Fig. 3A-B). The applied load (P) is related to the penetration depth of indenter (*δ*) as

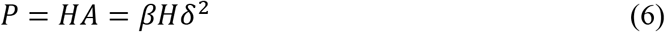

**Figure 3.**
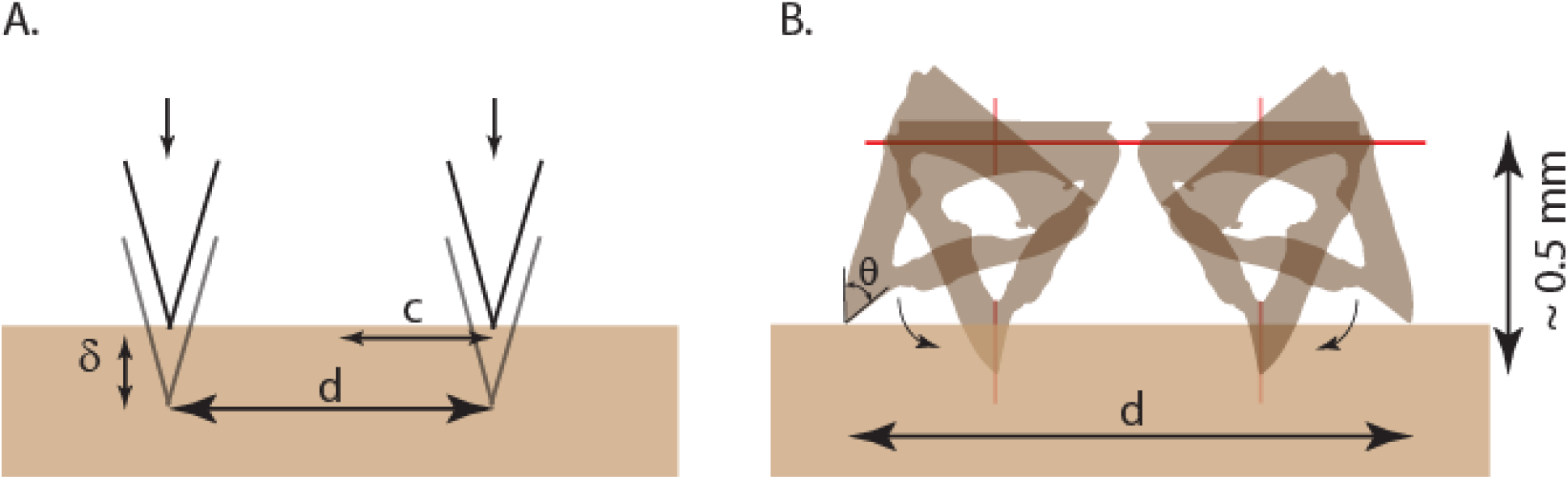
**A:** Schematic of the cutting mechanism modeled using the mandibles as two indenters penetrating the wood surface **B:** the corresponding possible mandible movement during cutting. The distance of axis of mandible movement from the tip is approximately 0.5 mm which is about the mandible height.

A is the projection of the contact area due to indentation and *β* is a constant which is dependent on the tip geometry. We consider an axisymmetric profile for the mandible, represented as an indenter, and given by a power law with exponent *γ*:

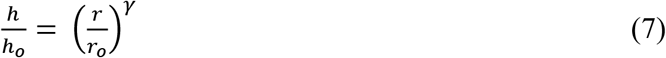

*r* is the radius and *h* is the height at a given location, and *r_o_, h_o_* are base radius and base height. Using the above relation, we can write the projected area of contact *A* during indentation as:

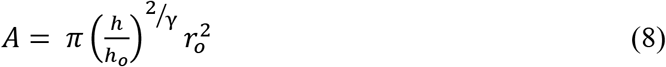

and inserting equations 5, 6, and 8 in equation 4, we write a general scaling law which describes the optimal chipping of the material as

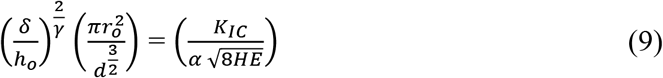

Known material parameters of the wood substrate occur on the right side of the equation whereas the distance between mandibles and the penetration depth of cut are on the left hand side of the expression. These equations also suggest that the animal may modulate the penetration depth of the indenter to achieve optimal chip formation for a given substrate. The wedge angle (θ) of the tip represents the vertex angle of the mandible was measured using the scanning electron micrographs of the mandible cross-section (Fig.3B).

## 3. Results and Discussion

### 3.1. Mandible morphology and zinc enrichment in the tip regions

The mandibles were visibly darker than the larval body (Fig. 1B). Scanning electron microscopy (SEM) characterization of the mandible morphology revealed a curved and sharp edge located towards the tip of the cutting regions with a relatively thicker base (Fig. 4A, B). The radius of curvature of the cutting edge, as measured in the cross-section of the polished mandible, was less than 0.5 μm (Fig. 4D); this is also characteristic of tools such as scalpel blades which typically have 0.25 μm. Energy dispersive X-ray spectroscopy detector (EDX) in the SEM, was used to characterize the presence of elemental transition metals, including zinc, at different spatial locations in the mandible (Fig. 5A). Analyses of the mandible surface shows a clear overlap of the peaks corresponding to zinc located preferentially in the tip region of the mandible but not in the base regions (Fig. 5B). Linear scans of the EDX spectra from cross-sections of the polished mandibles also show that the presence of zinc steeply increased in the tip region as compared to the base (Fig. 5C). The presence of zinc is hence exclusive to the cutting regions of the mandible tip whereas the neighboring regions are devoid of zinc. In an earlier study, we showed the importance of zinc in the ovipositor of fig wasp parasitoid insects (*Apocrypta* spp.) in increasing the cuticular hardness to aid insects in cutting through hard substrates [9]. A higher material hardness of ant mandibles is also known to correlate with presence of transition metals in the articulating surfaces [4]. We work on the hypothesis that zinc recruitment increases the hardness of the mandible tip as compared to base regions which do not come into contact with the wood substrate.

**Figure 4.**
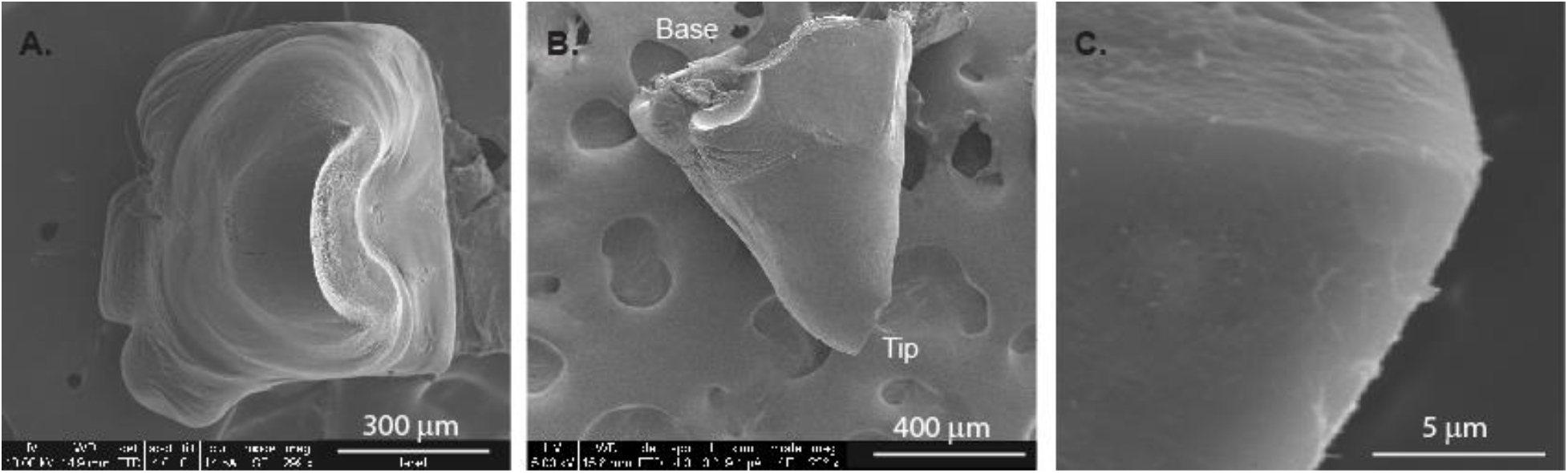
**A:** Scanning electron micrograph of the top of the CWSB mandible sample brings into focus the curved cutting parts of the tip region of the mandible. **B:** Side view highlights the surface of the mandible and the cutting cuspal edge. **C:** Zoomed view of the edge region of the mandible to demonstrate the tip sharpness which was measured to be 0.5 μm.

**Figure 5.**
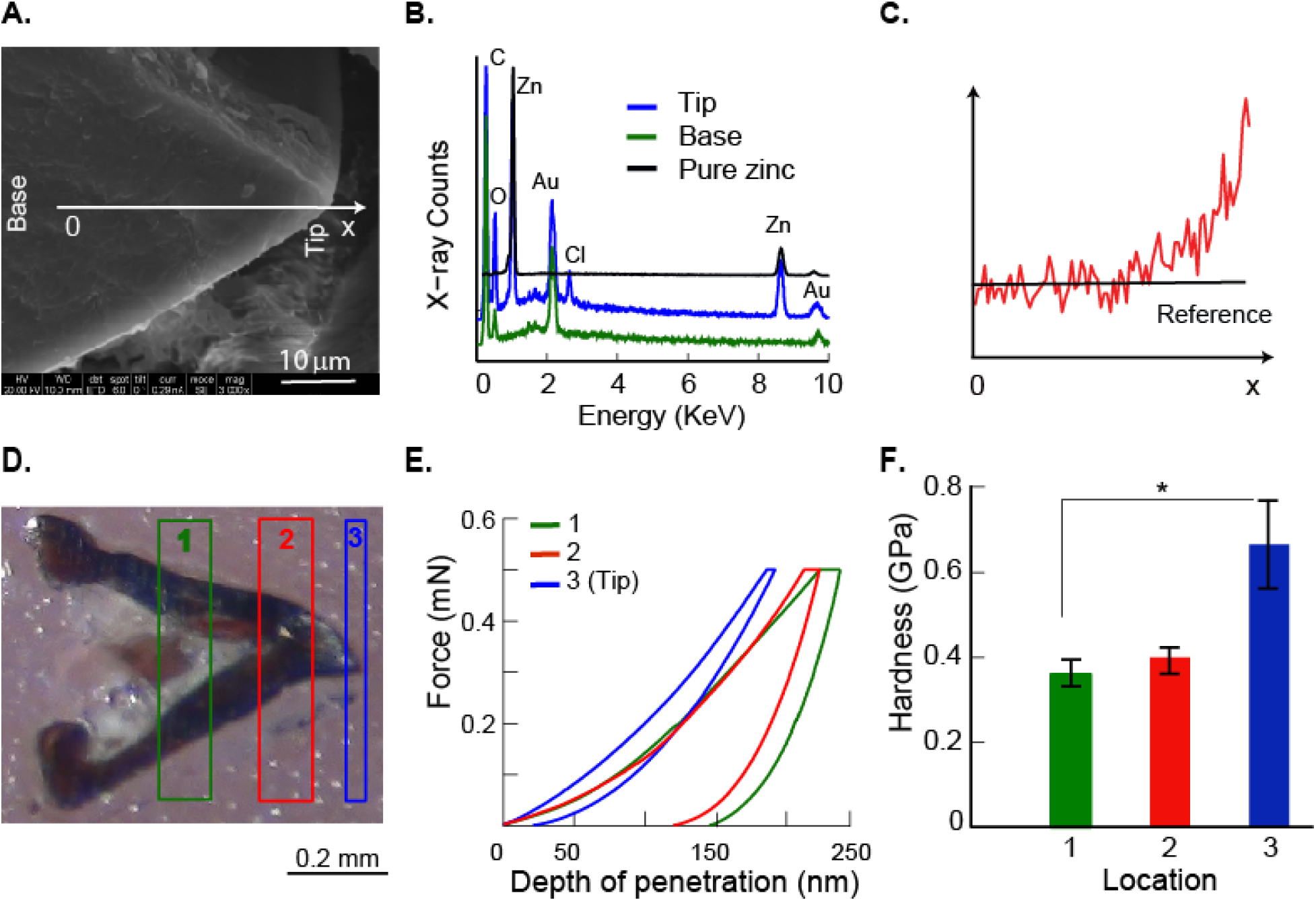
**A:** SEM image shows the base and tip regions of the mandible. **B:** Representative EDX spectra show peaks corresponding to zinc in the tip region alone as compared to the base. Related peaks from gold, chlorine, carbon and oxygen are also marked. **C:** Spectrum for a representative line scan from the base to tip in the specimen mandible shows increase in zinc content towards the mandible tip region as compared to the base. **D:** Three regions were selected for nanoindentation studies on a polished mandible sample to quantify possible differences in regional material properties. **E:** Force–depth relationships show lower values of final depth from region 3 (tip) as compared to other regions shown in 5D. **F:** Bar plots from indentation experiments show significantly higher hardness (p<0.05), plotted as average ± standard deviation, for the tip as compared to the base region.

### 3.2. Mechanical properties of the mandible and coffee wood

The force-depth curves, obtained using nanoindentation with a Berkovich indenter, showed clear differences in the mechanical properties from the tip of the mandible region as compared to other regions (Fig. 5E). We analyzed the force-depth relationships based on the standard Oliver-Pharr method in nanoindentation, and tested for variance in mechanical data between the tip, intermediate, and base regions of the mandible [27]. Our results showed that the contact stiffness in the intermediate (8.63±1.66 N/mm) and base (9.14±1.97 N/mm) regions of the mandible were similar (Table 1; Fig. 5F) and significantly higher as compared to the tip region (7.13± 0.87 N/mm; p<0.05). In contrast, mandible hardness in the tip region (0.67± 0.20 GPa; p<0.05) was significantly greater than both the base (0.36±0.06 GPa) and the intermediate region (0.40±0.06 GPa; Fig. 5F). These results are in agreement with earlier studies on incorporation of transition metals such as zinc, manganese, copper, and iron in the cuticle that show improved-material wear resistance by through increase in the local hardness [8]. The mandibles of termites (Cryptotermes primus) which feed on drywood were observed to be more scratch resistant as compared to the sub-families without the zinc and this was attributed to the presence of zinc [28].

**Table 1:** Stiffness and hardness from the three different mandible regions, marked in Fig. 5D, were calculated by analyzing the force-depth curves using nanoindentation results at each location (n=9) on a given specimen. Data are reported as average ± stdev for each sample; the last row in the table shows the overall average for each of the three regions.

| Sample | Stiffness (N/mm) |  |  | Hardness (GPa) |  |  |
| --- | --- | --- | --- | --- | --- | --- |
|  | Region1 | Region2 | Region3 | Region1 | Region2 | Region3 |
| 1 | 5.56±1.06 | 10.82±3.04 | 7.63±1.09 | 0.40±0.01 | 0.51±0.18 | 0.52±0.17 |
| 2 | 11.27±3.12 | 8.77±0.55 | 7.00±0.47 | 0.29±0.06 | 0.39±0.09 | 0.88±0.31 |
| 3 | 11.26±1.2 | 5.68±0.11 | 6.73±0.29 | 0.33±0.04 | 0.30±0.01 | 0.50±0.08 |
| 4 | 9.73±1.98 | 10.26±0.49 | 7.31±0.74 | 0.38±0.05 | 0.36±0.04 | 0.51±0.13 |
| 5 | 10.92±1.45 | 10.06±0.16 | 6.63±0.19 | 0.34±0.03 | 0.41±0.01 | 0.52±0.14 |
| 6 | 9.85±3.10 | 8.82±0.34 | 5.60±0.34 | 0.29±0.03 | 0.43±0.01 | 0.68±0.10 |
| 7 | 8.35±1.72 | 7.13±0.14 | 6.67±0.28 | 0.33±0.04 | 0.39±0.02 | 0.69±0.11 |
| 8 | 8.01±0.80 | 8.77±0.07 | 8.09±1.03 | 0.49±0.10 | 0.46±0.20 | 1.08±0.28 |
| 9 | 7.32±0.64 | 7.37±1.71 | 8.50±0.91 | 0.39±0.05 | 0.36±0.09 | 0.61±0.12 |
| **Average** | **9.14±1.97** | **8.63±1.66** | **7.13±0.87** | **0.36±0.06** | **0.40±0.06** | **0.67±0.20** |

Broomell and co-workers demonstrated the role of transition metals (Zn, Cu and Mn) in altering the hardness and stiffness of a model marine polychaete annelid jaw (*Nereis virens*) using differential chelation experiments [15,16]. These studies showed that the presence of transition metals, mediated through the crosslinking of histidine-rich proteins in the cuticle, resulted in a step-gradient in hardness. Such gradients in material properties along the mandible lead to redistribution in the stress distributions near the sharp contact and an improved ability in resisting crack propagation between two materially distinct regions through deflection [29]. The insect cuticle is a functionally graded composite material with hierarchical architecture. Finite element model of a spider fang, based on the differential organization of individual materials in combination with functional grading, demonstrate a structure which is adapted for damage resilience and effective structural stiffness [30]. How do the material properties of the mandible compare with that of wood? We used monotonic compression and micro-indentation experiments to quantify the mechanical properties of *Arabica* wood. The compression modulus of *Arabica* was 811±93 MPa. Hardness of wood samples was significantly lower (30.9 ± 5.3 MPa) as compared to that of the mandible tip (0.67± 0.20 GPa; p<0.01).

### 3.3. Finite Element Analysis of the mandible during a simulated bite

Are there any morphological adaptations in the mandible that make the larva effective in wood boring from a biomechanical perspective? X-ray microtomographic (XRadia Versa XRM 400) three dimensional reconstruction of the mandible revealed a highly curved and sharp cutting edge with an internally localized conical hollow region which is clearly visible in the sagittal (Fig. 6A-C, Movie S2), transverse and coronal projections (Fig. 6D). Volume of the hollow region was approximately half the total mandible volume (53.0 ± 8.3 %, n=3; AMIRA 6.0.1, FEI software) which makes the mandible a relatively light structure (Movie S2). We investigated the role of this hollow region in regard to the stress distributions on the mandible during substrate boring. We developed a finite element model as discussed in the Methods section and included graded material properties from the nanoindentation studies on the discretized mandible structure (ABAQUS v6.12). Three different boundary conditions were applied which are referred as fully pinned, semi pinned and pivot (Fig. 2D). The fully pinned condition allowed us to investigate the load bearing capability of the mandible. The semi pinned case includes the effects of abductor and adductor attachments. Finally, the pivot case takes into account displacement of the mandible about the hinge axis (M-M’) which is caused by the actuation of muscles at the base of the mandible.

**Figure 6.**
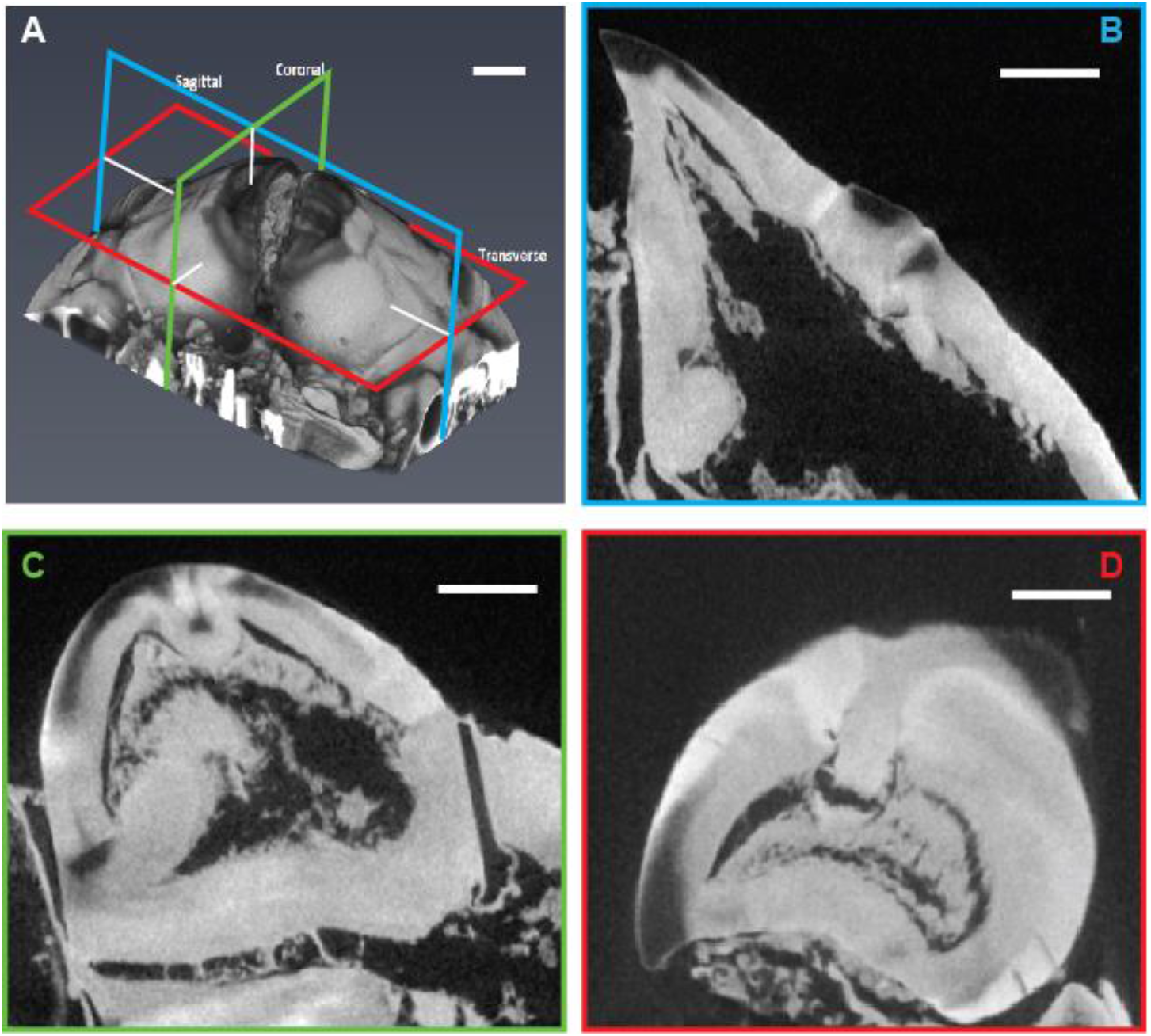
**A:** Three dimensional rendered microCT image of a borer mandible with highlighted sectional planes. **B:** Projection along sagittal plane shows a conical hollow part and a sharp cutting edge. **C:** Projections along coronal plane and **D:** transverse plane shows curved mandibular region with a bean shaped relatively hollow region.

Figure 7 shows stress contours on various surfaces of the mandible for three different boundary conditions. Few differences are visible in the stress distributions at the tip of the mandible (region 3) which is primarily involved in cutting; a maximum stress of ~ 180 MPa was present on back surface. Stress differences among the different boundary conditions are apparent towards the muscle attachment points (region 1) and at the base surface of the mandible. To explore for possible spatial variations, we plotted stresses along the front and back outer bounding surfaces of the mandible on the sagittal section T-T’ (Fig. 2D). The start of the hollow region within the mandible is highlighted in gray in these figures. These results show that stresses had similar profiles for the fully pinned and semi pinned cases on the front surface of the mandible (Fig. 8). Stresses were ~4.5 times higher in Region 1 located towards the base of the mandible for the pivot case as compared to the two other cases. Stresses on the back surface decreased monotonically after a maximum traction in the intermediate region; stress profiles were similar in the tip and intermediate regions for all three cases for the back surface (Fig. 8B). Because of the concentrated reaction force provided by surface S2, the semi pinned case had the highest stress in Region1 (Fig. 2D).

**Figure 7:**
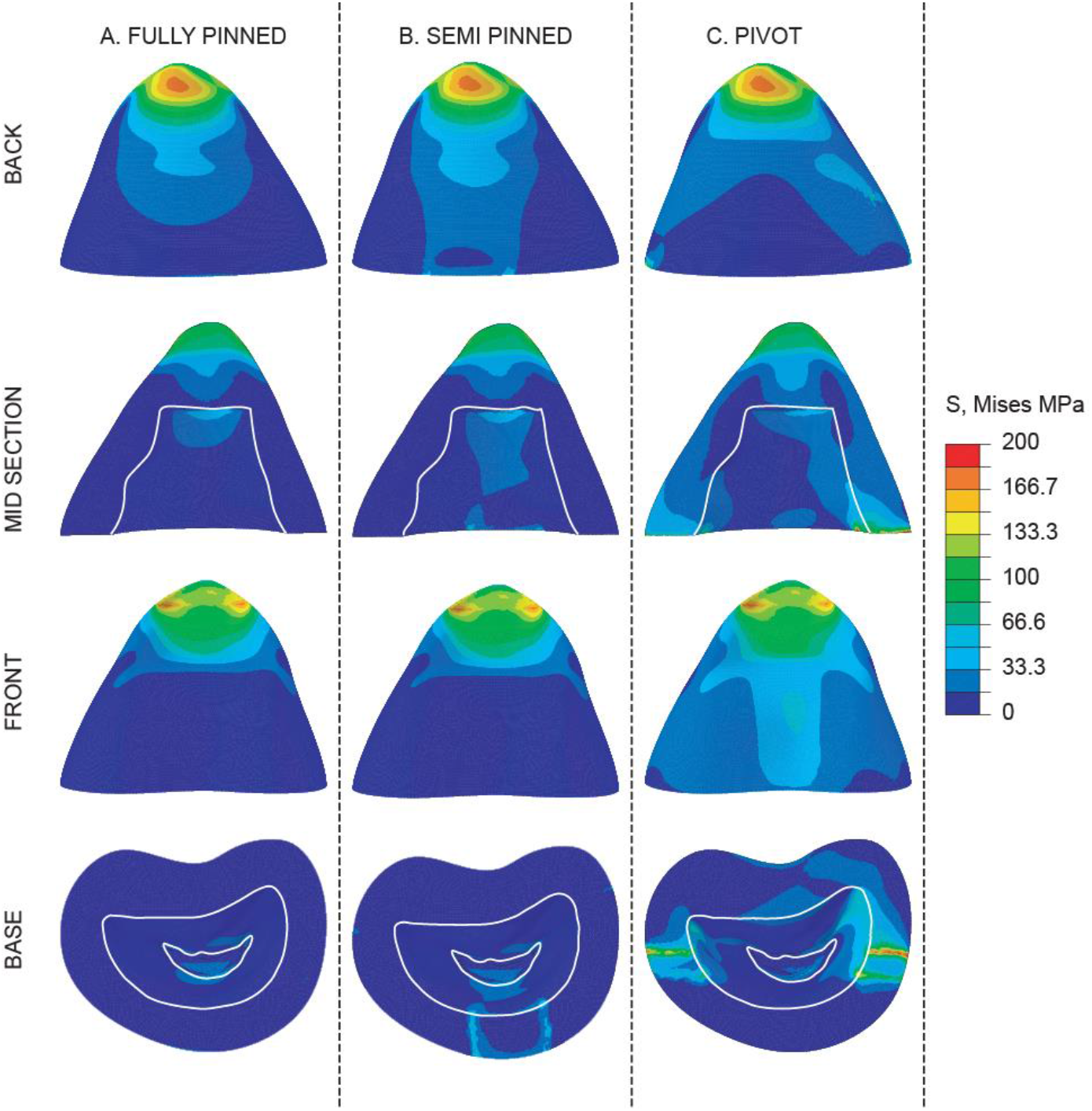
von Mises stress distributions corresponding to A) fully pinned, B) Semi pinned, and C) Pivot boundary conditions are shown on the back, mid-section, front, and base surfaces of the mandible.

**Figure 8:**
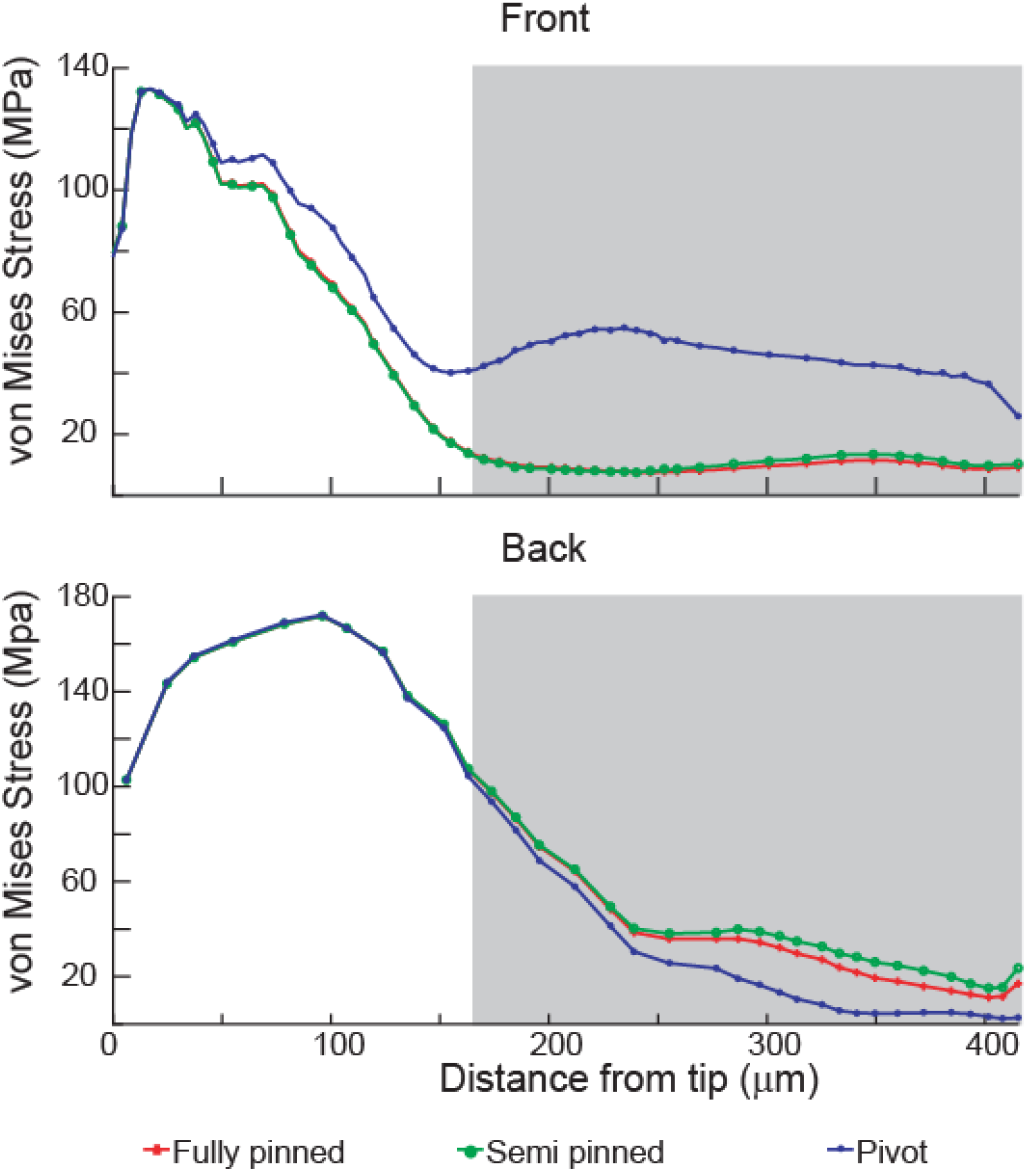
Comparison of stress distributions on the **A)** front and **B)** back surfaces of the mandible along the sagittal section (T-T’) shown in Figure 2D. The distributions are plotted for the three different boundary conditions which were simulated in the study starting from the tip region.

The hollow structure in the mandible extended from ~165 microns from tip to the base of mandible. This span coincides with regions on the back and front surfaces having less stress (<90 MPa). The midsection surfaces in all the three simulations (Fig 7) show that in locations near the hollow region, marked by the white boundary, there is sudden drop in stresses on both the front and back surfaces. Recent studies on ants which tunnel through wood showed the presence of short, cone-shaped mandibles with wide articulatory base which rotate and help with concentrating muscle-generated forces to zinc-reinforced mandibular tips [22]. A similar rotator hinge mechanism with long mandibular output levers operate in male stag beetles to achieve forceful bites [23]. The presence of a hollow region also increases the toughness in absorbing the loads during impact loads which is similar to the design of antlers that present the highest toughness for mineralized materials [31]. These biomechanical adaptations may hence aid in the cutting of hard wood by mandibles.

### 3.4. Penetration into wood through mandibular action

Mechanical design of the mandibles hence reveals a combination of novel morphological and structural principles that make the beetle extremely effective in boring through wood. The low aspect ratio (~1.25) in mandible structures hence enhances their stiffness. Similar aspect ratios (1.45) are found in scorpion chela that also withstands high stress when feeding on hard shell preys [32]. The loading conditions in our simulations were idealized using solutions proposed for a cantilever beam of variable cross-sectional area [33]. Bending stress for a cantilevered structure, σb = M/Z, is low at the base where moment of inertia (I) and Z are large. In contrast, σb is high near the tip regions which experiences uniform surface tractions during wood cutting (Fig. 7A-C). Hollow structures are also known to be stiffer than solid structures for a given material volume due to the greater moment of inertia and have lower material volume without compromising the function [5]. The mandible shape also determines the maximum size of the substrate chip which is an essential functional requirement in ingesting wood [34].

We developed a general scaling law to quantify the penetration of the mandible into wood using a fracture mechanics framework (Fig. 3A, 3B). The formation of wood chips may hence be controlled by the beetle through modulation of *δ* which is given by the expression.

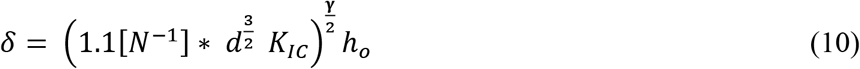

K_Ic_ in this expression is the mode I fracture toughness of coffee wood. Because of unavailability of the value of K_Ic_ we used an average value corresponding to Douglas fir 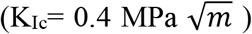 based on available literature [35]. Using the microCT data, we estimated the biting distance between mandibles to be between 0.1 and 0.5 mm. Equation (10) shows that the depth of cut is hence proportional to the distance between mandibles during biting. We used average measured mechanical properties of the coffee wood measured from microindentation experiments (H=30.9 MPa and E=1.6 GPa) to calculate the penetration depth during cutting of the wood substrate by the mandibles using the scaling model. We also explored variations in the indenter shape, *γ*, with the distance between indenters to estimate the values of *δ* using the model (Figure 9A-C). The values of *δ* for ~100 μm distance between mandible tips are shown for variations in γ values from 0.97-1.0 under the assumption that the penetration depth cannot exceed the mandible height (Table 2). To explore the dependence of the depth of optimal cut on *K_Ic_*, we parametrically varied values over a range of values (Table 3). A small vertex angle may result in tool failure in tool design whereas a high vertex angle reduces the cutting ability of the tool [36]. By parametrically varying the vertex angles from 64° to 82° in turning processes showed that a wedge tool with 64° vertex angle was the most efficient for material removal [37]. The angle measurements from the cross-section of mandible tip were in the range of ~47-52 ^o^ when approximating it as a simple wedge. Scanning electron micrographs of the bored regions of Arabica wood show pattern (Fig. 10A, B) that show wood damage caused by the mandibles. These data show that loads applied on the wood fibers using a specialized tool helps produce small chip sizes which are suitable for ingestion without the tool itself undergoing fracture during cutting. The shape of the mandible edge seems to be optimized for optimal cutting of wood chips. Physiological responses including detoxification of defensive allele-chemicals and breakdown of secondary metabolites with resident microbes also aid the beetle in the ingestion of wood [38].

**Figure 9.**
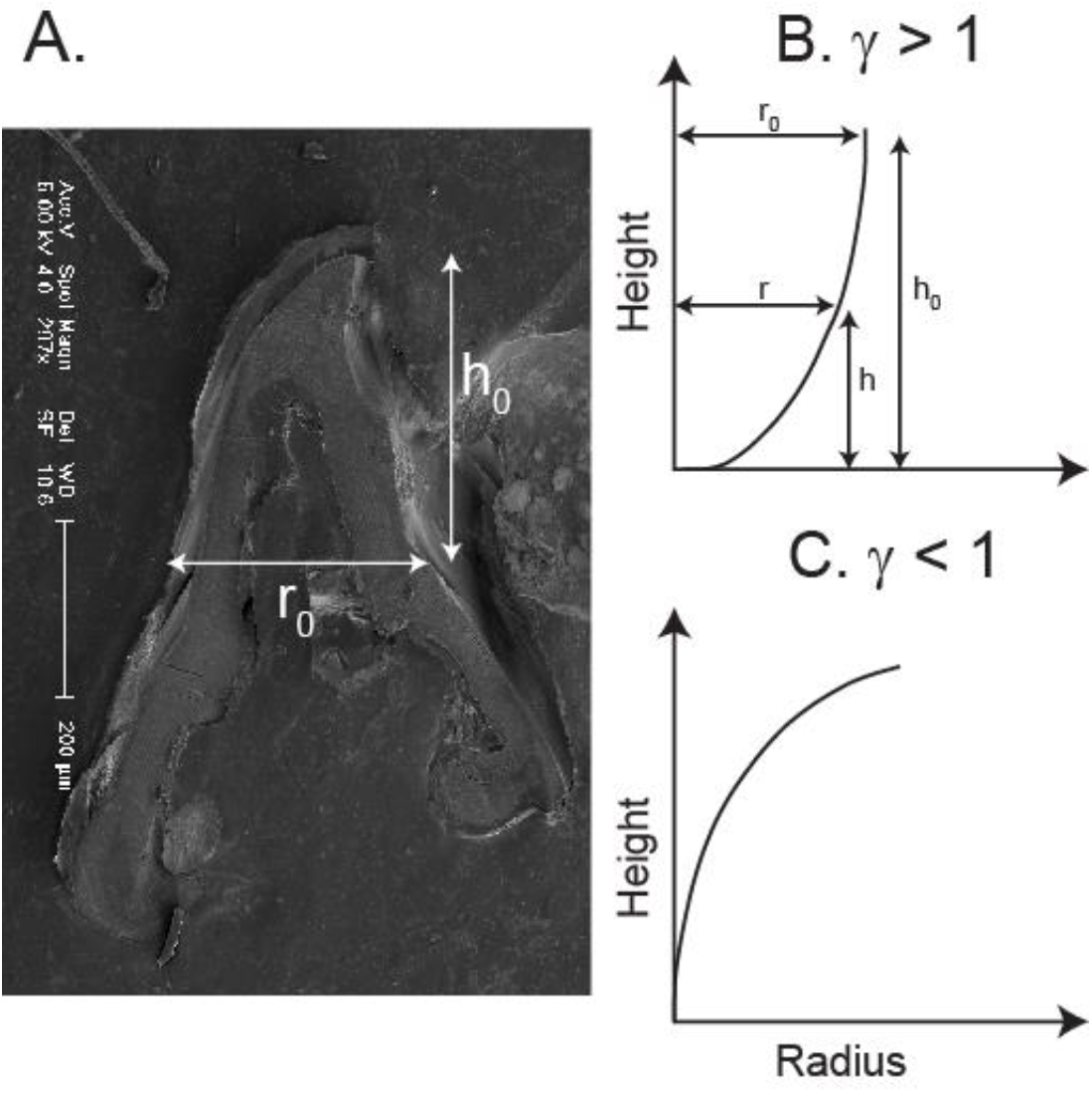
**A:** SEM image of the mandible with representative diameter and height marked on the image. Schematic representations of the mandible model, assuming symmetric half edges, show variations in the radius (*r*) with height (*h*) for two cases with different shape factor values corresponding to **B:** *γ*> 1 and **C:** *γ*<1. *r_o_, h_o_* are base radius and base height.

**Figure 10.**
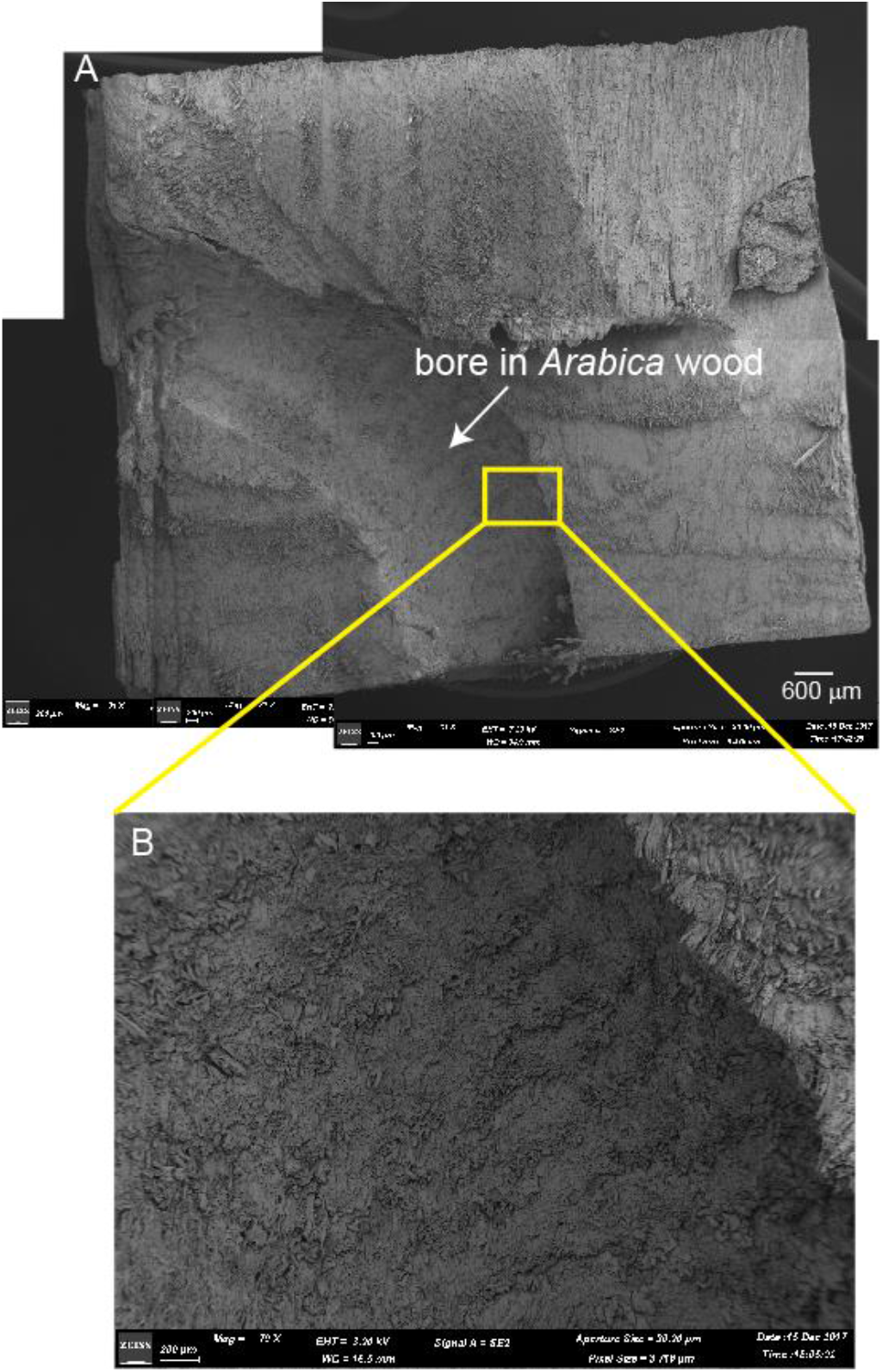
**A:** SEM of the *Arabica* wood sample shows the bored region which was obtained after sectioning the sample. **B.** Enlarged bored region shows a pattern of fractured wood fibrils that was caused by the sharp mandible tip.

**Table 2:** Depth of cut value, *δ*, was estimated by varying the shape parameter, γ, and using two different values of separation between indenters, d, with 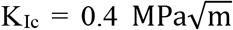 corresponding to Douglas fir (26). Predicted values of *δ* are within the range of plausible values with the assumption that depth of cut is of the order of the height of the mandible.

| **d**<br>($\mu\text{m}$) | $\gamma$ | $\delta$<br>($\mu\text{m}$) | | **d**<br>($\mu\text{m}$) | $\gamma$ | $\delta$<br>($\mu\text{m}$) |
| --- | --- | --- | --- | --- | --- | --- |
| 500 | 0.01 | 444 |  | 100 | 0.01 | 438 |
| 500 | 0.1 | 476 |  | 100 | 0.1 | 422 |
| 500 | 0.5 | 655 |  | 100 | 0.5 | 358 |
| 500 | 0.8 | 831 |  | 100 | 0.8 | 316 |
| 500 | 1 | 974 |  | 100 | 1 | 291 |
| 500 | 1.2 | 1140 |  | 100 | 1.2 | 268 |
| 500 | 1.4 | 1340 |  | 100 | 1.4 | 247 |
| 500 | 1.5 | 1450 |  | 100 | 1.5 | 237 |
| 500 | 2 | 2160 |  | 100 | 2 | 193 |

**Table 3.** Depth of cut, δ, was estimated by varying K_Ic_ for two different cases with variations in the bite distance, d. δ varies directly with K_Ic_ under conditions for the cracks to meet and create a chip. A variety of chip sizes may be produced for a given value of *γ* by varying the distance between the mandibles and the penetration depth.

| **d**<br>**($\mu\text{m}$)** | **$\gamma$** | **$\delta$**<br>**($\mu\text{m}$)** |  | **d**<br>**($\mu\text{m}$)** | **$\gamma$** | **$\delta$**<br>**($\mu\text{m}$)** |
| --- | --- | --- | --- | --- | --- | --- |
| $K_{IC}=0.04 \text{ MPa}\sqrt{\text{m}}$ | | | | $K_{IC}=4 \text{ MPa}\sqrt{\text{m}}$ | | |
| 100 | 0.8 | 125 |  | 100 | 0.8 | 794 |
| 100 | 1 | 92 |  | 100 | 1 | 920 |
| 100 | 1.2 | 67 |  | 100 | 1.2 | 1067 |
| 500 | 0.8 | 330 |  | 500 | 0.8 | 2086 |
| 500 | 1 | 307 |  | 500 | 1 | 3079 |
| 500 | 1.2 | 286 |  | 500 | 1.2 | 4544 |

## 4. Conclusions

There are several key implications of this study. First, from a biomechanics perspective we show a highly sharp structure used to initiate fractures in wood, graded material properties in the mandible structure, peak stresses in the mandible tip and reduced bending stresses at the mandible base, and lower material volume that does not compromise its function during cutting. Incorporation of zinc in the cutting edges alone increases the material hardness; this facilitates ease in cutting through hard wood. Zinc enrichment in larval mandibles suggests advantages during developmental to increase the material hardness. Finally, the shape of the mandible was compared to that of standard indenters and the tip angle in the mandible was similar to wedge tools. We developed a scaling law by modeling the interaction of the two mandibles with a quasi-brittle like mechanical behavior of the wood substrate and show that the larvae can produce small chip sizes suitable for ingestion through modulation of the distance between the two mandible cutting edges depending on the wood properties. These adaptations make the beetle a remarkably efficient and stealthy animal that eludes detection; this makes it a successful pest in economically important crops like coffee. These unique design principles may also be extended into other comparative systems, such as red palm weevils in coconut palm, Asian long head beetles in maples etc., which face similar challenges in cutting of hard substrate. Biomechanical studies may aid in identifying zinc chelators as herbicides to control infestation and also brings into focus mechanical principles in nature to design bioinspired tools.

## Acknowledgments

We thank Mr. Akash Vardhan for help with Amira for image reconstructions and Prof. Sanjay Sane for useful discussions. We are grateful to Mr. Joshua Amirtharaj, Tata Coffee Ltd, for providing CWSB samples and for discussions. NG acknowledges Department of Science and Technology for the Ramanujan fellowship and research grant that supported part of this work and to the Central Coffee Research Institute for project support. NMP is supported by the European Commission under the Graphene Flagship Core2 No. 785219 (WP14 “Polymer Composites”) and FET Proactive “Neurofibres” grant No. 732344. N.M.P and L.K. are supported by Fondazione Caritro under “Self-Cleaning Glasses” No. 2016.0278.

